# High Quality Phasing Using Linked-Read Whole Genome Sequencing of Patient Cohorts Informs Genetic Understanding of Complex Traits

**DOI:** 10.1101/2022.03.28.486092

**Authors:** Scott Mastromatteo, Angela Chen, Jiafen Gong, Fan Lin, Bhooma Thiruvahindrapuram, Wilson WL Sung, Joe Whitney, Zhuozhi Wang, Rohan V Patel, Katherine Keenan, Anat Halevy, Naim Panjwani, Julie Avolio, Cheng Wang, Guillaume Côté-Maurais, Stéphanie Bégin, Damien Adam, Emmanuelle Brochiero, Candice Bjornson, Mark Chilvers, April Price, Michael Parkins, Richard van Wylick, Dimas Mateos-Corral, Daniel Hughes, Mary Jane Smith, Nancy Morrison, Elizabeth Tullis, Anne L Stephenson, Pearce Wilcox, Bradley S Quon, Winnie M Leung, Melinda Solomon, Lei Sun, Felix Ratjen, Lisa J Strug

## Abstract

Phasing of heterozygous alleles is critical for interpretation of *cis*-effects of disease-relevant variation. For population studies, phase is often inferred from external data but read-based phasing approaches that span long genomic distances would be more accurate because they enable both genotype and phase to be obtained from a single dataset. To demonstrate how read-based phasing can provide functional insights, we sequenced 477 individuals with Cystic Fibrosis (CF) using linked-read sequencing. We benchmark read-based phasing with different short- and long-read sequencing technologies, prioritize linked-read technology as the most informative and produce a benchmark phase call set from reference sample HG002 for the community. The 477 samples display an average phase block N50 of 4.39 Mb. We use these samples to construct a graph representation of *CFTR* haplotypes, which facilitates understanding of complex CF alleles. Fine-mapping and phasing of the chr7q35 trypsinogen locus associated with CF meconium ileus demonstrates a 20 kb deletion and a *PRSS2* missense variant p.Thr8Ile (rs62473563) independently contribute to meconium ileus risk (p=0.0028, p=0.011, respectively) and are *PRSS2* pancreas eQTLs (p=9.5e-7 and p=1.4e-4, respectively), explaining the mechanism by which these polymorphisms contribute to CF. Phase enables access to haplotypes that can be used for genome graph or reference panel construction, identification of *cis*-effects, and for understanding disease associated loci. The phase information from linked-reads provides a causal explanation for variation at a CF-relevant locus which also has implications for the genetic basis of non-CF pancreatitis to which this locus has been reported to contribute.

## Introduction

Current genetic epidemiological studies often fail to capture the complete diploid nature of the human genome (1) largely because of a reliance on genotyping arrays and short-read whole genome sequencing (WGS). These technologies can identify heterozygous alleles but provide little to no information regarding the *cis* or *trans* phase relationships of their heterozygous allele pairs. Accurate haplotype information can be essential in informing phenotype-genotype relationships. One of the most well-known examples come from compound heterozygosity in monogenic disorders such as cystic fibrosis (CF) (2).

CF is caused by mutations in the CF transmembrane conductance regulator (*CFTR*) (3). Over 2,100 variants have been identified in *CFTR* (4), greater than 400 of these have been shown to be disease causing while others are reported to have varying clinical consequence and are CF-causing only when in *cis* with another deleterious variant (5). Meanwhile, individuals with identical CF-causing alleles display variable disease severity and response to *CFTR*-targeting therapies (6). CF co-morbidities and variation in disease severity are complex genetic traits (7), presumed to be due to the impact of other genes beyond *CFTR*, referred to as modifier genes. For example, genome-wide association studies (GWAS) of CF meconium ileus (MI), an intestinal obstruction seen at birth in 13–21% of individuals with CF (8), have identified associated loci (9)(10)

By design, the GWAS arrays mostly contain common SNPs in easily accessible regions of the genome. The MI associated SNPs do not appear in high linkage disequilibrium (LD) with protein coding variations, suggesting their impact is through gene regulation. However, much remains to be learned about the variation that is in *cis* with these associated SNP risk alleles or whether combinations of multiple *cis*-acting variants contribute to MI risk; for this, genotype data at the associated loci must be phased.

In a typical epidemiological study, data external to the target individual is used to reconstruct maternal and paternal haplotypes. Pedigree-based phasing offers a high degree of accuracy (11) but requires a family-based experimental design and cannot resolve phase for variants that are heterozygous for all members. Population-based phasing is a cost-effective alternative that exploits shared ancestry information and linkage disequilibrium (LD) patterns to statistically infer haplotypes. However, the statistical nature of population-based phasing makes it vulnerable to frequent switch errors: accidental transitions from maternal to paternal haplotypes between neighbouring heterozygous sites (1). Phasing rare variants can also be problematic, requiring inference when few or no copies of that rare variant are present within the reference population.

In contrast, individual-level phasing approaches determine phase relationships for a target individual without reliance on an external dataset. Sequencing reads that overlap multiple heterozygous sites are phase informative (12) but the maximum phase distance is restricted by the size of the sequencing read which makes short-read data ineffective when attempting to phase over non-trivial distances. Long-read sequencing technologies such as Pacific Biosciences (PacBio) SMRT sequencing and Oxford Nanopore generate longer reads capable of phasing longer distances, but these technologies are often error-prone, very costly or both. Other alternatives utilize a standard short-read sequencing pipeline with an additional experimental step that introduces long-range information into the read data. For example, the 10x Genomics (10XG) linked-read technology (13) and Universal Sequencing Technologies TELL-Seq (14) tag reads derived from a single DNA molecule with a shared nucleotide barcode, enabling otherwise independent reads to be linked and capable of long-distance phasing.

Here, we benchmark the phasing capabilities of different sequencing technologies using public data from the well-studied individual NA12878 with the practical goal of scaling to sample sizes large enough to quantify haplotype distributions for statistical analysis. This work guides our choice of 10XG as a technology to apply to the Canadian CF Gene Modifier Study Consortium (CGMS) cohort. We sequenced 477 individuals with CF from the CGMS using 10XG linked-read technology at approximately 30x coverage and leverage the data to improve understanding of a MI associated locus.

MI GWAS has identified three genome-wide significant loci and a suggestive intergenic locus within the T-cell receptor beta region (chr7q35) (9) that was replicated in independent samples (10). Early sequencing work in the chr7q35 region identified five trypsinogen paralogs with approximately 90-91% nucleotide similarity which were annotated as T4 to T8 (15). Cationic trypsinogen *PRSS1* (T4) and anionic trypsinogen *PRSS2* (T8) are major forms of trypsin found in the pancreas, one of the earliest affected organs in CF (16). The other three genes are pseudogenes: *PRSS3P1* (T5), *PRSS3P2* (T6) and *TRY7* (T7); of the three, there is only evidence for *PRSS3P2* transcription but no known evidence of a protein product (17). The GRCh38 reference genome only include three of these genes (T4, T5, T8) which is an accurate representation of a common deletion polymorphism that removes T6 and T7. This approximately 20 kb deletion appears to have arisen via non-allelic homologous recombination (18) and this represents a common variation found in approximately 41% of individuals with European ancestry (19). The GRCh38 alternative contig, KI270803.1 (20), represents the non-deleted haplotype and contains genes T4-T8. This is further complicated by reference assembly GRCh37 being erroneously structured (T4, T5, T6) and excluding *PRSS2*; a correction was later released (chr7_gl582971_fix) that included all five genes.

In the present study, we provide a phasing benchmark using different technologies, summarize the phasing quality achievable across the 477 individuals with CF and use the phase information to unravel the complex genomic architecture at the chr7q35 modifier locus.

## Results

### Comparison of phasing potential between read technologies

Here we consider the phasing quality of four different sequencing technologies: 10XG linked-reads, PacBio continuous long-reads (CLR), PacBio circular consensus sequence (CCS; branded as HiFi), and Nanopore reads. Phased variant calls for reference individual NA12878 is assessed for each technology (data sources listed in Supplementary Table 1). Variant calls fall into discrete phase blocks: a set of variants that are phased with respect to each other. Nanopore and 10XG technologies demonstrate longer, more contiguous phase blocks than PacBio CLR or CCS (Figure 1a). Phase blocks for chromosomes 1-3 are shown in Supplementary Figure 1 and additional statistics are available in Supplementary Table 2.

**Fig. 1.**
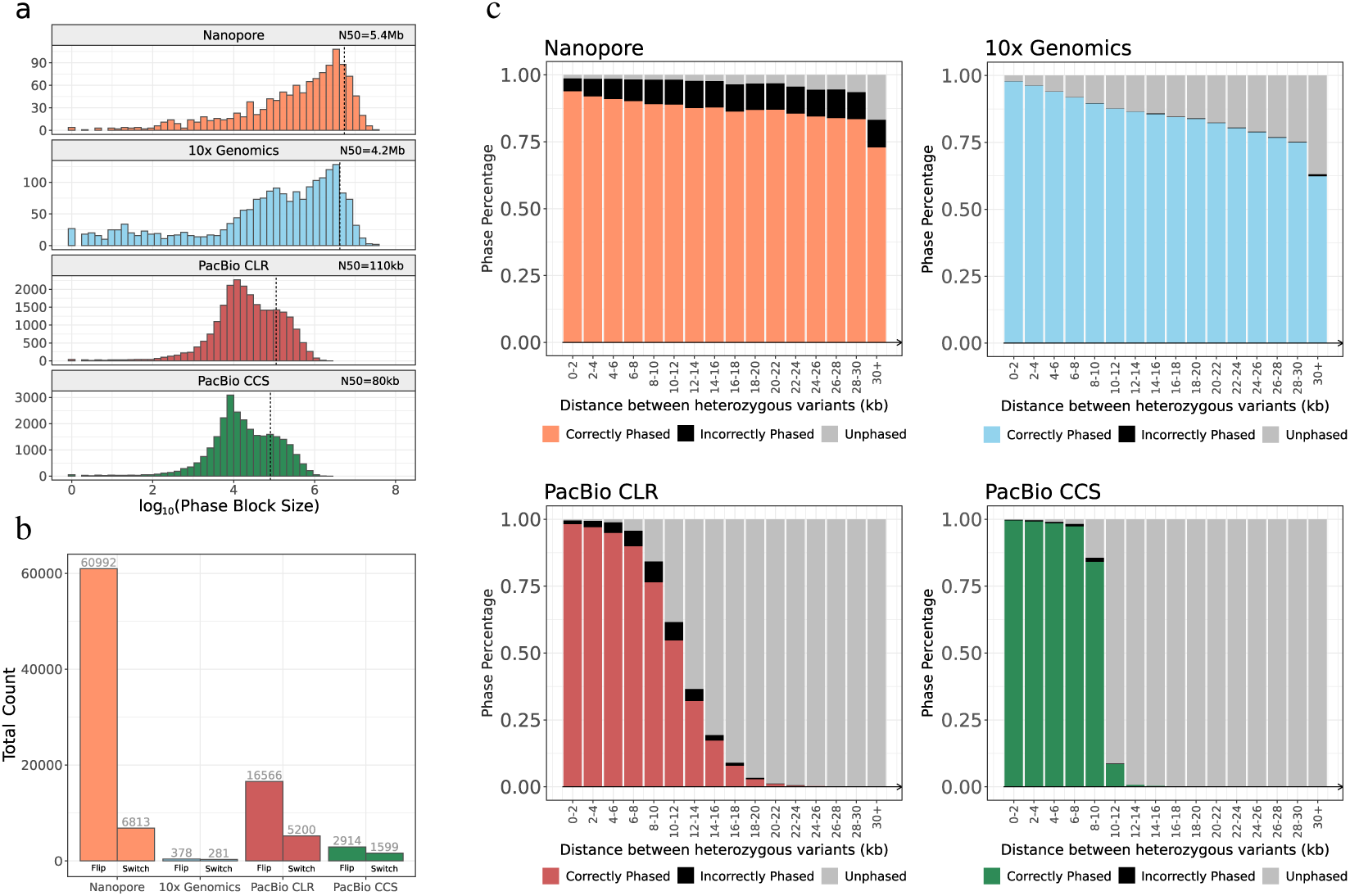
Benchmarking phased calls using four different sequencing technologies against NA12878. Phased VCFs produced for reference individual NA12878 using Nanopore, 10XG, PacBio CLR and PacBio CCS. **a** Distribution of phase block lengths for each technology. N50 is annotated as dotted line. Nanopore: 1231 blocks, N50=5.41 Mb; 10XG: 2021 blocks, N50=4.19 Mb; PacBio CLR: 24129 blocks, N50=110 kb; PacBio CCS: 27407 blocks, N50=80 kb. **b** Number of flip and switch errors identified by comparison to Platinum Genomes benchmark using pairs of heterozygous variants. **c** Phase accuracy shown as a function of the distance between variant pairs in kb. A variant pair is deemed correctly phased if the phase configuration agrees with the Platinum Genome truth set; a variant pair could also disagree with the truth set (black) or have no phase call made (grey).

We assess phase accuracy by comparison with the NA12878 Platinum Genomes phase calls generated from a 17-member pedigree (21). Figure 1b shows disagreements with the benchmark as flip errors (a single variant on the wrong haplotype) and switch errors (a contiguous block of variants on the wrong haplotype). Nanopore demonstrates the lowest accuracy with 97.28% of assessed variant pairs agreeing with the benchmark. PacBio CLR and CCS show higher accuracy (99.12% and 99.82%, respectively) but 10XG has the best performance with 99.97% accuracy and only 659 flip and switch errors total across all assessed variant pairs. The phasing accuracy of PacBio and 10XG specifically has been previously reported (22).

The relationship between phase quality and the distance between pairs of neighbouring variants is presented in Figure 1c. Nanopore and 10XG technologies are capable of phasing variant pairs spanning tens of kilobases, correctly phasing 75% and 84% of adjacent heterozygous variants in the range of 28-30 kb, respectively. In contrast, the CLR and CCS PacBio data show a significant drop-off in the ability to phase heterozygous variant pairs that are greater than 10 kb apart, corresponding to the expected read lengths of these technologies. PacBio CLR technology correctly phases 2% of the variants within the 10-20 kb range compared to 2.7% for CCS.

The accuracy of short-reads in conjunction with the long-range information offered by 10XG linked-reads creates long phase blocks while maintaining a low error rate relative to the other technologies. Although Nanopore reads generate a more contiguous set of phase blocks, it comes at a cost of a higher error rate. It is critical to minimize incorrect phase calls because even a single switch error produces a multitude of misleading pairwise variant relationships by splitting a consistent phase block into two completely out-of-phase parts. With this consideration, 10XG produces the highest quality phase calls of the technologies assessed.

The insight into the strengths and weaknesses of each technology motivates an approach to combine multiple technologies and improve phase quality. We devised a pipeline to combine phased phased variant call format (VCF) files from multiple sources to generate a consensus phase set and benchmark the results against the NA12878 Platinum Genomes truth set. Supplementary Figure 2 shows the effect different combinations of sequencing technologies has on phase properties. Using all four technologies in combination, 99.93% of variants are phased in 894 phase blocks (N50=13.24 Mb) with an accuracy of 99.59% (complete statistics in Supplementary Table 3).

We apply this consensus phasing strategy to create a high-quality phased VCF for the well-studied individual HG002. The Genome in a Bottle (GIAB) Consortium provides phased benchmark calls for small variants in HG002 (23) generated from parental genotypes and Strand-seq data. However, many phasing errors can be detected in the GIAB release VCF by manual assessment of the read data (Supplementary Figure 3). In release version 4.2, 28.2% of heterozygous variants remain unphased in the release VCF. To improve this resource, we generate a consensus by combining the following data: Strand-Seq, 10XG linked-reads, PacBio CCS, PacBio CLR, Nanopore and include the pedigree phase information available in the GIAB release VCF for HG002. The consensus of the six sources of phase information phases 99.996% of heterozygous variants within 81 phase blocks with N50 of 90.3 Mb across the entire genome. This data is available at (24).

### Phasing 477 Canadians with cystic fibrosis

We performed whole genome sequencing of DNA for 477 individuals from the CGMS cohort (10) using the 10XG linked-read technology at 30x depth (25x after trimming the 10XG barcode). The phasing distance of 10XG linked-reads is limited by the size of DNA molecules extracted. We investigated different extraction methods and found MagAttract produces the best results, consistent with the publicly available NA12878 sample (Figure 2a-d). Mean molecular length averages 58.7 kb (range: 32.6-95.4 kb) across 463 MagAttract extracted samples and is a strong predictor of the quality of the phasing. The average MagAttract extracted sample is phased in 2444 blocks, with N50 of 4.39 Mb and a mean of 1428 variants per block. The largest phase block across all samples is 247.97 Mb and all but two samples have >97% of all genes shorter than 100 kb phased in a single block. Additional statistics can be found in Supplementary Table 4.

**Fig. 2.**
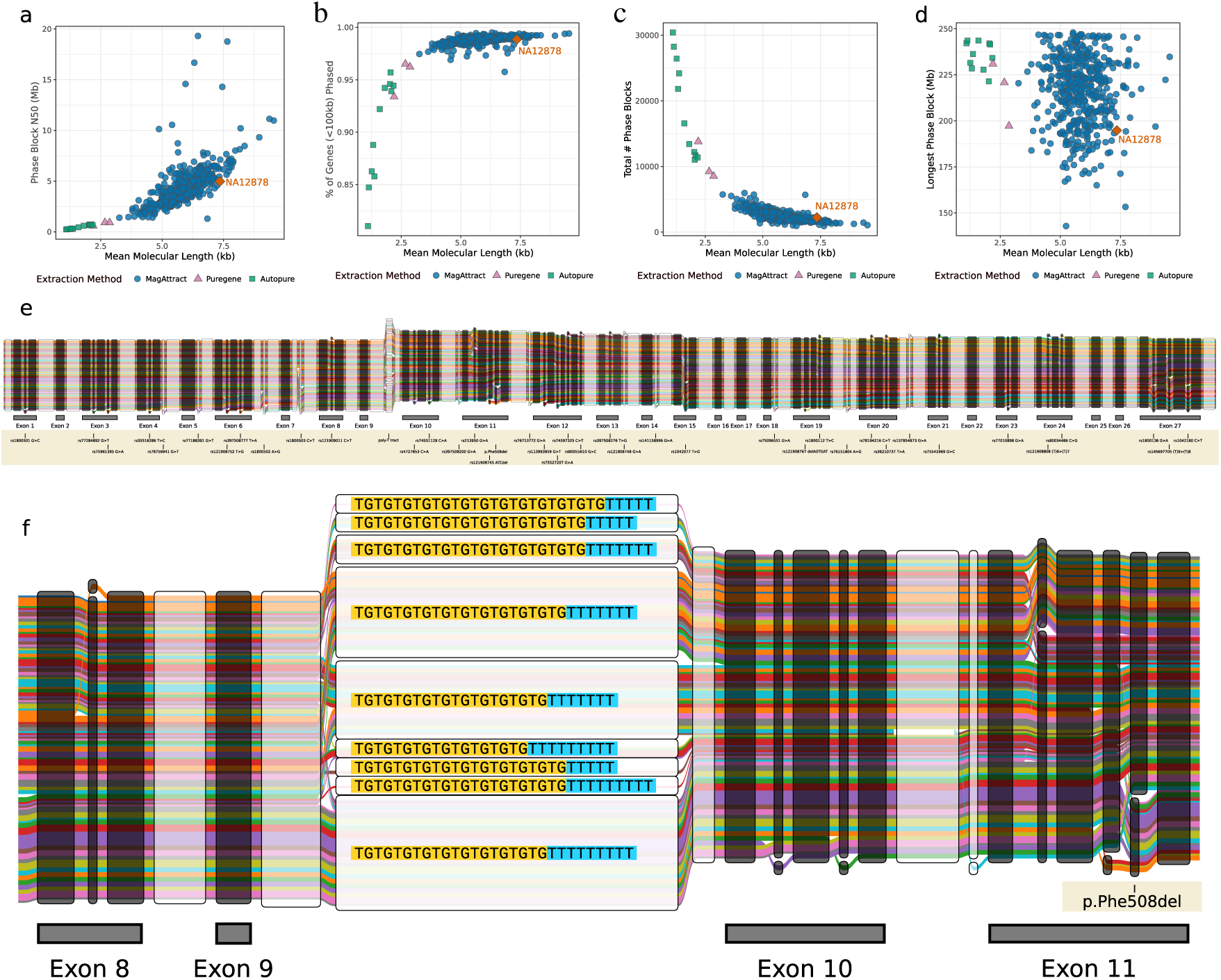
Genome-wide phasing statistics versus mean molecular length for CGMS samples and NA12878 sequenced by 10XG. DNA from CGMS cohort extracted using either MagAttract (blue circle), Autopure (green square) or Puregene (purple triangle). GIAB data for NA12878 (orange diamond) was down-sampled to a comparable coverage (30x). Statistics are compared against mean molecular length reported by Long Ranger (13) **a** Phase block N50; **b** Proportion of phased genes with length less than 100 kb; **c** Total number of phase blocks; **d** Size of longest phase block in base pairs. **e** Graph representation of exonic variants for 898 *CFTR* haplotypes. The graph is composed of nodes representing sequence and haplotype groups as colored edges. The complete haplotype sequence can be reconstructed by concatenating the nodes along a path. The thickness of each edge denotes the haplotype frequency in the dataset. Nodes belonging to exons are annotated and colored black. **f** The intronic poly-T tract is included in the graph representation. Nine different poly-T alleles are visualized here and shown with respect to three SNPs downstream from the poly-T tract.

To complement genome-wide statistics, we assess the local phasing of a 389 kb region encompassing the CF causal gene, *CFTR* (GRCh38 chr7:117379963-117768665; *CFTR* plus 100 kb on both sides). The most common CF-causing variant is p.Phe508del; 241 individuals homozygous for this variant comprise about half of the sequenced samples. Due to a conserved haplotype, individuals homozygous for p.Phe508del possess high levels of homozygosity along the entire *CFTR* gene which makes it difficult to phase. The median p.Phe508del homozygous individual has 10 heterozygous variant calls within the assessed region (one per ∼40 kb) compared to 236 heterozygous variants (one per ∼1.6kb) for the median individual with heterozygous CF-causing alleles. Consequently, 152 of the 199 individuals with heterozygous CF-causing variants have a single phase block spanning the complete 389kb region. This demonstrates how the phasing of causal loci in disease cohorts with a recessive mode of inheritance could pose unique challenges for read-based phasing techniques but also highlights the potential to identify complex alleles that may explain disease variation (7).

We construct a graph representation of the phased sequence at the *CFTR* locus from 449 individuals with CF to provide a visual understanding of the 10XG-derived haplotypes (Figure 2e). The graph includes the multiallelic poly-T tract polymorphism to highlight how a graph representation of haplotypes can inform disease phenotypes. Variation at the poly-T tract results in altered splicing and can cause CF if in *cis* with specific *CFTR* mutations (25); p.R117H in phase with a short poly-T is CF-causing while the clinical manifestations for those with longer poly-T sequence is less certain. Nine different poly-T alleles are visualized and their phase is shown with respect to downstream variants including p.Phe508del.

### Analyzing the chr7q35 trypsinogen CF modifier locus

The MI GWAS-suggestive locus on chr7q35 has five duplicated trypsinogen paralogs (*PRSS1, PRSS3P1, PRSS3P2, TRY7* and *PRSS2*) but is structurally variable across reference assemblies (Figure 3a). The GRCh38 chr7 sequence includes a large deletion polymorphism that removes *PRSS3P2* and *TRY7*. Reads from individuals who carry a non-deleted haplotype align spuriously to GRCh38, resulting in false variant calls (Supplementary Figure 4). Realignment of 10XG reads to alternative contig KI270803.1 improves the calling and phasing of variation and enables the large deletion polymorphism to be unambiguously called (Supplementary Figure 5). Among individuals with European ancestry, we find almost no variation within the deletion boundary on haplotypes lacking the deletion (Supplementary Figure 6). A simple genotype coding of the deletion sufficiently captures the genetic variation contained in this subregion and is used for all subsequent analysis.

**Fig. 3.**
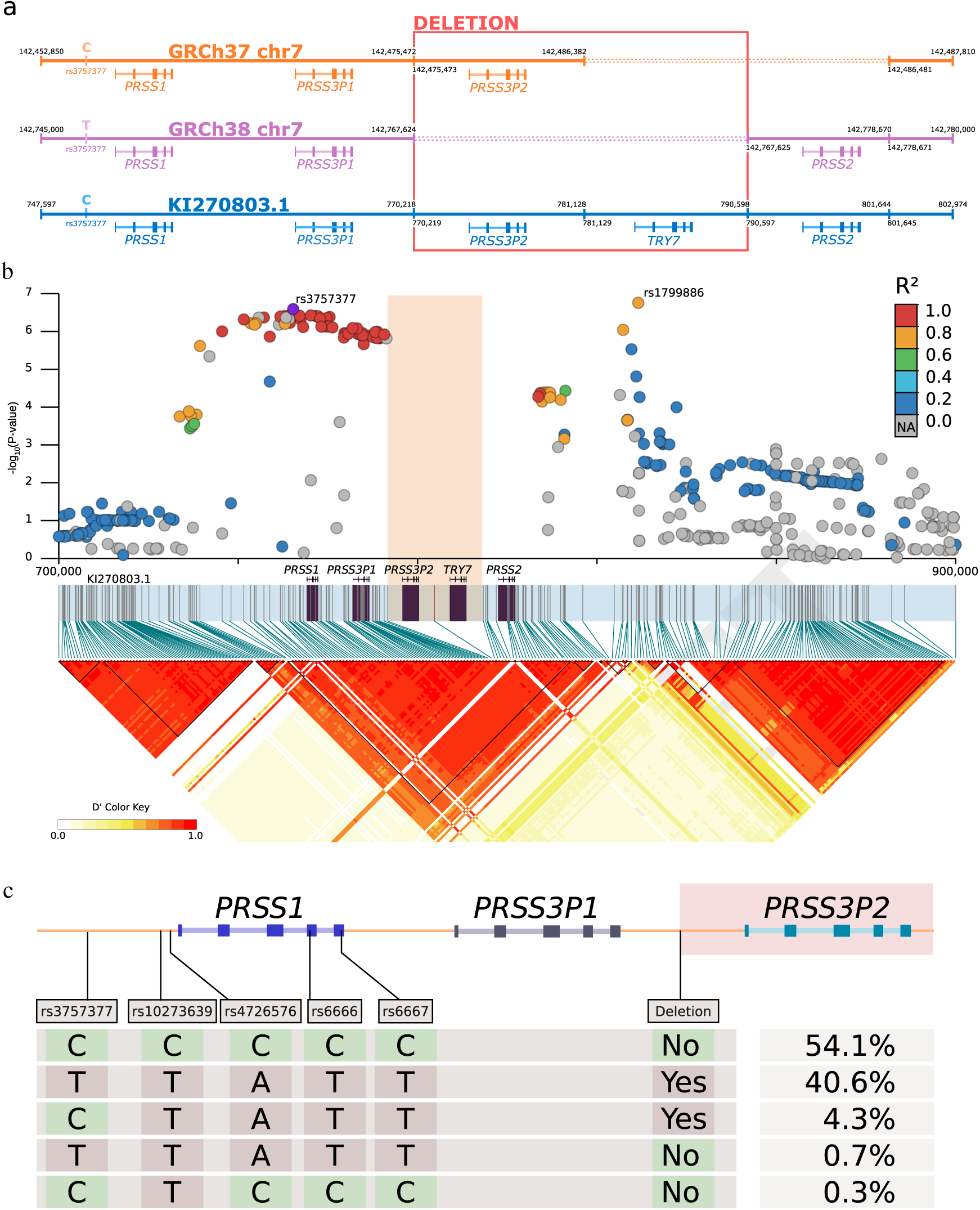
Characterizing the chr7q35 trypsinogen locus. **a** Differences between chromosome 7 reference assemblies for GRCh37, GRCh38 and alternative contig KI270803.1. In the GRCh37 assembly, *TRY7* and *PRSS2* are absent. The GRCh38 assembly does not include *PRSS3P2* and *TRY7* because it accurately represents a haplotype with a common ∼20 kb deletion polymorphism (highlighted in red). KI270803.1 represents a haplotype without the deletion polymorphism. **b** LD matrix calculated from 10XG phased calls; deletion allele is denoted by orange rectangle. Haplotype blocks are drawn as black triangles, all five trypsinogen homologs are located within a single block (KI270803.1:737033-802909). MI GWAS summary statistics lifted from GRCh37 to KI270803.1 are shown, r2 with respect to rs3757377. **c** Four SNPs in the same LD block as rs3757377, phased with the deletion polymorphism. SNPs include a common pancreatitis risk allele (rs10273639) (26), a *PRSS1* promoter SNP (rs4726576) that alters expression of a reporter gene in mice (27), two synonymous *PRSS1* variants (rs6666 and rs6667). Five unique haplotypes are observed in 10XG data, the frequencies are shown as a percentage. The two major haplotypes account for 94.7% of the observed data.

424 of 477 10XG samples are completely phased in a single block across a conservative 200 kb region surrounding the *PRSS1*-*PRSS2* locus (KI270803.1:700000-900000). This phase information elucidates the LD structure of this locus for the CGMS cohort and is shown alongside MI GWAS summary statistics in Figure 3b. Two association peaks centered at rs3757377 (KI270803.1:750284C>T) and rs1799886 (KI270803.1:823812T>C) are present in different LD blocks. The rs3757377 risk allele “T” has a frequency of 41% in the 10XG calls. We phase this SNP with respect to other variants of interest within the same LD block, the two major haplotypes account for 94.7% of the observed data (Figure 3c).

The second peak centered at rs1799886 has a similar minor allele frequency of 43.5% but is not in strong LD with the deletion polymorphism (D’=-0.55, r^2^=0.19). A search for variants in *cis* with rs1799886 reveals a nonsynonymous PRSS2 variant (p.Thr8Ile), rs62473563 (KI270803.1:793978C>T), with 10.7% minor allele frequency and a high D’ with rs1799886 (D’=-0.98, r^2^=0.09). The rs1799886 “T” allele is in *cis* with p.Thr8Ile for 100 out of 101 haplotypes. The GWAS signal is tagging this protein-coding SNP; this relationship was not uncovered in the original analysis of the GWAS results due to the absence of PRSS2 from the GRCh37 reference.

A query of the Genotype-Tissue Expression (GTEx) v8 data (28) was conducted to search for pancreas eQTLs with respect to the five trypsinogen paralogs. *PRSS3P2* and *TRY7* are not reported by GTEx v8 due to their absence from the GRCh38. *PRSS3P1* does not have significant pancreas eQTLs but this is expected as it is not transcribed. Significant pancreas eQTLs are reported for *PRSS2* (Supplementary Table 5) but not for *PRSS1*. This result is surprising because there is a common SNP in the promoter region that is reported to alter *PRSS1* expression (27) but did not appear as a significant eQTL. LocusFocus (29) detects colocalization between MI association p-values and GTEx v8 *PRSS2* pancreas eQTLs (colocalization p-value=7.1e-8, Supplementary Figure 7). This suggests that MI risk could be modulated by altered *PRSS2* expression. The reliability of these results depends on accurate accounting of the 20 kb deletion polymorphism during read alignment to GRCh38. We found that the presence of the extra 20 kb sequence did not significantly alter the normalized gene expression counts for *PRSS1* or *PRSS2* when compared with GTEx v8 counts (r^2^ correlation of the two datasets >0.99, Supplementary Figure 8).

To improve comparison to the predominantly European CGMS data, 252 GTEx samples with the race labelled as “white” were used to recalculate pancreas eQTLs. The GTEx v8 variant calls for these samples were lifted to KI270803.1 and the deletion polymorphism was imputed using the 10XG CGMS samples as a reference panel. Similar to the GTEx v8 results, there are no significant (p<0.05) eQTLs for *PRSS1* (Supplementary Figure 9) but *PRSS2* has pancreas eQTLs (Figure 4a). The imputed deletion polymorphism appears as a strong *PRSS2* pancreas eQTL (p=7.8e-5). Conditioning on the deletion polymorphism reveals that rs62473563 (*PRSS2* missense variant, p.Thr8Ile) acts as an independent eQTL (Figure 4b). Conditioning on rs62473563 increases the significance of the deletion polymorphism (Figure 4c) and conditioning on both eliminates the *PRSS2* eQTL signal (Figure 4d). The presence of p.Thr8Ile and the deletion polymorphism are both associated with reduced *PRSS2* expression. This conditional analysis is summarized in Table 1.

**Fig. 4.**
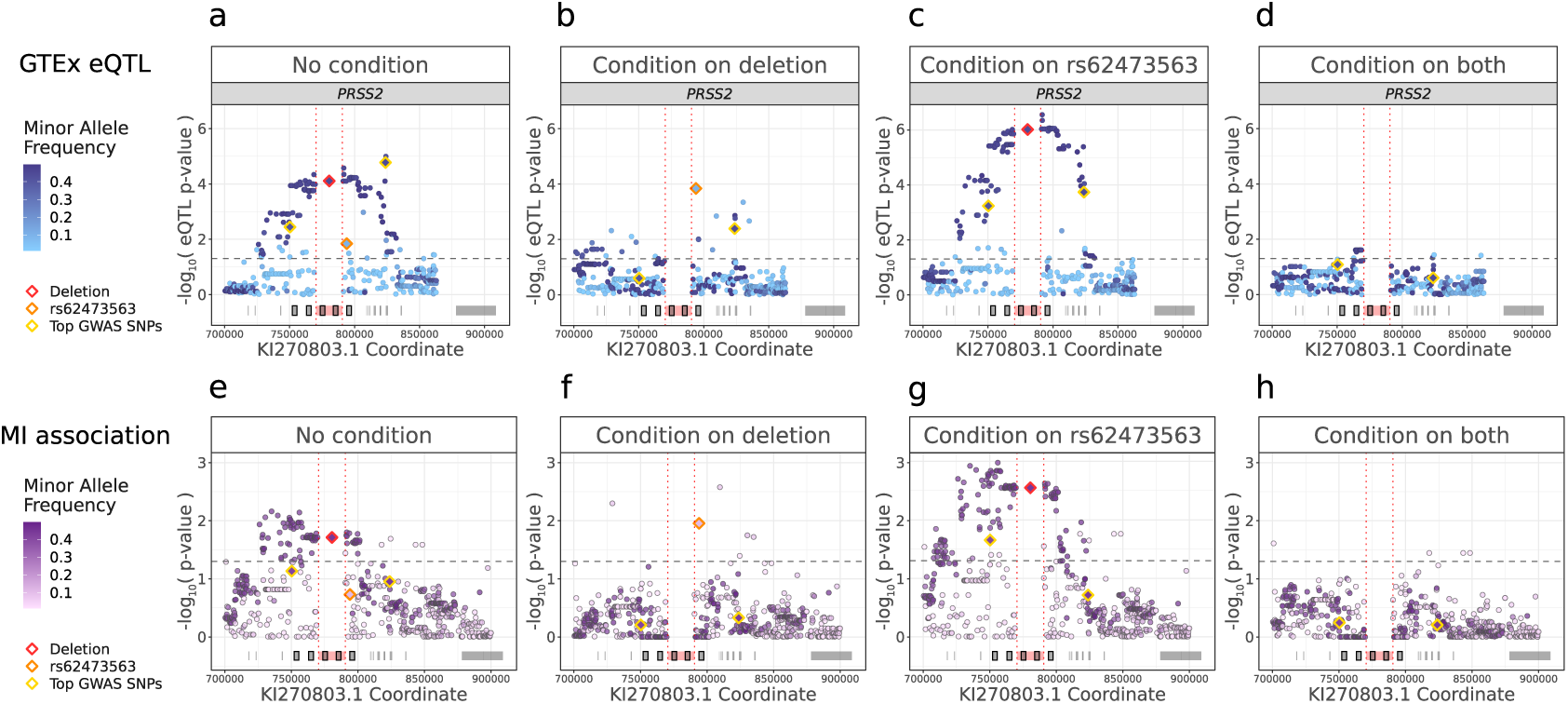
Conditional association analysis reveals common association pattern for *PRSS2* pancreas eQTLs and MI risk. GTEx v8 variant calls lifted to KI270803.1 and deletion allele (red diamond) imputed from 10XG calls. **a** Recalculated *PRSS2* pancreas eQTLs. **b** GTEx *PRSS2* pancreas eQTLs conditioning on deletion polymorphism. **c** *PRSS2* pancreas eQTLs conditioning on rs62473563 (orange diamond). **d** GTEx *PRSS2* pancreas eQTLs conditioning on both rs62473563 and deletion polymorphism. **e** Association with MI was similarly performed for 309 10XG samples. **f** MI risk conditioning on deletion polymorphism. **g** MI risk conditioning on rs62473563. **h** MI risk conditioning on both rs62473563 and deletion polymorphism.

**Table 1.** GTEx *PRSS2* pancreas eQTL analysis and association to MI risk using 10XG data under conditional analysis.

| Variant | Conditioned on | *PRSS2* pancreas eQTL<br>n=252 |  | MI association<br>n=309 |  |
| --- | --- | --- | --- | --- | --- |
|  |  | Slope (SE) | P-value | Beta (SE) | P-value |
| rs62473563 | - | -0.24 (0.10) | 0.014 | 0.42 (0.32) | 0.19 |
| rs62473563 | Deletion | -0.38 (0.097) | 1.4e-4 | 0.93 (0.37) | 0.011 |
| Deletion | - | -0.24 (0.060) | 7.8e-5 | 0.53 (0.22) | 0.019 |
| Deletion | rs62473563 | -0.31 (0.060) | 9.5e-7 | 0.75 (0.25) | 0.0028 |

To understand these eQTL results in the context of MI, we analyzed genotyping array data from CGMS cohort individuals (n=2635, Supplementary Figure 10). The deletion polymorphism is associated with an additive increased risk of disease (beta=0.29, p=5.2e-4) but imputation for rs62473563 is poor. Instead, we performed fine-mapping using the 10XG sequencing calls for whom MI status and 10XG data were available. A similar association pattern observed for the pancreas eQTL was recapitulated for the MI phenotype in 337 individuals sequenced with the 10XG technology. (Supplementary Figure 11). Interestingly, the contribution of this locus in CF individuals with two minimal function *CFTR* alleles appears attenuated, which is likely due to their already elevated risk due to *CFTR* (8). Exclusion of 28 individuals with minimal function CF alleles produces stronger evidence of association with MI despite the smaller sample size (Figure 4e-h, Table 1). Notably, the *PRSS2* variant p.Thr8Ile remained associated with MI after accounting for the deletion polymorphism in the model (beta=0.93, p=0.011). Both deletion and p.Thr8Ile are associated with a reduction in *PRSS2* expression and a higher risk of MI.

## Discussion

Phasing of genetic sequence improves understanding of causal variation at GWAS-associated loci, especially in regions of complex genetic architecture and when allelic heterogeneity is present. However, haplotype reconstruction is typically not a priority when studying disease cohorts following-up GWAS identified loci. Here we demonstrate that linked-read technology provides a robust and cost-effective option for epidemiological studies of complex loci.

Benchmarking different read technology against Platinum Genomes highlights the exceptional phasing accuracy produced by 10XG linked-reads. Large phase blocks with N50 upwards of 4 Mb are achievable with this technology – more than sufficient for studying targeted loci. It should be noted that the Nanopore and PacBio data used in this study were based on public availability and technological improvements have been made since those datasets were released. The general insights offered by the benchmarking comparison still apply to newer iterations of these technologies.

While 10XG linked-reads provides high-quality phase information, we observed that the linked-reads often generated incomplete phase blocks where many variants remain unphased within a block. Variants with insufficient phase-informative reads occur stochastically, especially in positions with low coverage. Unphased variants can also be the result of regions with low mappability for short reads. In contrast, long-read technologies generate more uniform coverage, improve mappability and produce complete phase blocks. To achieve the most reliable phase calls for a single individual, we show that an ensemble approach can compensate for the individual deficits of each technology by taking a consensus of multiple callsets. We have produced and made available a consensus VCF for the well-studied GIAB sample HG002. This consensus is a useful reference for studies interested in benchmarking phase calls, since HG002 has one of the most well-studied genomes and, to our knowledge, has yet to be comprehensively phased.

To demonstrate the practical utility of phased sequence data for a cohort, we investigated the chr7q35 trypsinogen locus that did not reach genome-wide significance in our largest GWAS of MI in CF to date (10). Nonetheless this locus was tantalizing due to the role trypsinogen plays in digestion and the specificity to the pancreas, one of the organs most significantly impacted in CF. The architecture of the chr7q35 trypsinogen locus requires careful analytic consideration. The region is heavily susceptible to reference bias, where differences between which reference assembly is used can produce misleading results. Reference bias in this locus has had documented clinical consequences, specifically the detection of a pathogenic *PRSS1* variant called based on misaligned reads derived from trypsinogen pseudogenes (30)(31). We mitigated misalignments by using reference sequence KI270803.1 that provides a more complete representation of this locus. The reference bias issues here motivate the general need to transition from linear references to more comprehensive representations such as graph-based references that can capture and accommodate the range of variation found within a population. The construction of these graphs can also benefit from the read-based phasing made available through technologies such as linked-reads, as demonstrated by the *CFTR* graph we present here.

The chr7q35 trypsinogen locus, and *PRSS1* in particular, is well-studied in the context of non-CF pancreatitis. An amino acid substitution in *PRSS1* (p.R122H) is the most common cause of hereditary pancreatitis in Europeans (32). This small change alters a trypsin cleavage site that is important for regulation of trypsin activity through autoinactivation of trypsinogen (33). Similarly, chronic pancreatitis has been shown to be associated with a common T>C variant (rs10273639) near *PRSS1* (26), thought to be associated with altered risk by tagging a promotor SNP (rs4726576) that increases *PRSS1* expression (27). Increased genetic risk of pancreatitis is typically manifested as increased trypsin activity, by the production of more functional trypsin or greater resistance to degradation via autoinactivation (34). Despite the depth of evidence supporting a relationship between *PRSS1* and pancreatitis, there is not the same level of support for *PRSS2*. Transgenic human *PRSS2* in mice has been shown to aggravate pancreatitis (35) and the *PRSS2* variant p.G191R promotes degradation and provides some protection against chronic pancreatitis (36). This supports the hypothesis that PRSS2 activity may also contribute to pancreatitis risk.

The data presented here suggests a more relevant role for *PRSS2* over *PRSS1* in MI. We identify two putatively contributing polymorphisms that independently alter MI risk and *PRSS2* expression: a 20 kb deletion polymorphism and a non-synonymous variant in exon 1 of *PRSS2*. These polymorphisms are in *cis* with risk variants in two independent MI associated SNP clusters, confirming the evidence of allelic heterogeneity seen in our previous MI GWAS (10). The deletion polymorphism is in *cis* with the common SNP rs10273639 found to alter non-CF pancreatitis risk (26). While previous work has suggested a connection between this haplotype and *PRSS1* expression, the results presented in this current work do not implicate *PRSS1* expression as the mechanism. The association between rs10273639 and *PRSS1* expression was initially established using 69 pancreas tissue samples after removal of 3 outliers (33). However, the raw data shows positive correlation between *PRSS1* and *PRSS2* expression (r2=0.83) and suggestive evidence of an association between rs10273639 and *PRSS2* (p=0.053, Supplementary Figure 12). While the data was interpreted to support *PRSS1* expression as a causal explanation, it does not exclude a *PRSS2* contribution. Given the extreme transcriptional activity of this locus in pancreatic cells, it would not be surprising that a structural change caused by the large 20kb deletion polymorphism upstream of the *PRSS2* promoter could alter *PRSS2* transcription.

A second MI GWAS association signal is in near-perfect linkage with the p.Thr8Ile variant in *PRSS2* (rs62473563). When restricted to a European subset, this variant is also the most significant *PRSS2* pancreas eQTL. Conditioning on the deletion polymorphism, p.Thr8Ile also showed evidence of increased MI risk in the 10XG samples highlighting its independent effect. PRSS2 trypsin operates extracellularly and therefore must be targeted for the endoplasmic reticulum (ER) during translation. The first 15 amino acids contain the sequence specific for binding of the signal recognition particle (SRP) targeting for the ER. An amino acid change here can alter SRP recognition efficiency which triggers a translation quality control (37). As p.Thr8Ile is a common variant found in healthy individuals, it does not seem consequential enough to cause a disease phenotype in isolation, but perhaps it is sufficient to modify severity of phenotypes when found in combination with disease states such as CF.

Non-CF pancreatitis is related to increased trypsin activity, typically attributed to *PRSS1* (26). For MI we see the opposite relationship where more trypsin activity reduces risk, and our data suggests this is due to *PRSS2* expression variation. Although there is conflicting evidence of whether *PRSS1* or *PRSS2* is the relevant gene, in both contexts the haplotype with the common deletion polymorphism is associated with lower levels of trypsinogen. Similarly, the presence of p.Thr8Ile is associated with lower *PRSS2* expression and higher MI risk; the effect on non-CF pancreatitis – if any – has not been reported to our knowledge. As MI is a neonatal intestinal blockage caused by thick and adhesive consistency of the first stool, a simple explanation is that higher trypsin levels in the intestine break down and discourage the formation of this blockage-causing stool, thereby reducing risk. In fact, it is known that the meconium of individuals with CF contain high levels of protein (38) and more active trypsin could provide a protective effect against blockage.

## Conclusions

This study demonstrates the benefit of sequencing technologies that simultaneously informs genotype and phase for a given individual. Construction of phased haplotypes enables greater insight into *cis*-effects at complex loci. Additionally, insights made available though LD structure, genome graphs and reference panel construction are also dependent on phase information. Here we identify a 20 kb deletion polymorphism and *PRSS2* missense variant that alters risk of complex CF traits and is associated with *PRSS2* gene expression. This could not have been elucidated without the phase information made available through 10XG linked-reads. It was therefore discouraging to receive news during this study that 10x Genomics was discontinuing their linked-read sequencing with no intention to make it available through other providers. We hope analogous methods such as Universal Sequencing Technologies TELL-Seq and long-read technology such as PacBio SMRT sequencing and Oxford Nanopore continue to mature to allow the research community continued access to read-based phasing that is cost-effective for population studies. Technologies that capture phase information are paramount to a complete understanding of GWAS loci, contributing to a greater understanding of genetic epidemiology.

## Methods

### Retrieval and phasing of benchmark genomes

NA12878 benchmarking variant calls were downloaded from Illumina’s Platinum Genomes (version 2017-1.0) (21). High confidence variant calls for reference individual HG002 were downloaded from the Genome in a Bottle (GIAB) Consortium (version 3.3.2 and 4.1). Sequencing datasets for both individuals were collected from multiple platforms including 10XG linked-reads, PacBio Circular Consensus Sequence (CCS) and PacBio Continuous Long Reads (CLR) and Oxford Nanopore. Phased variant calls respect to the reference genome GRCh38 p.12 were either downloaded directly or aligned and phased. Direct links to each dataset are provided in Supplementary Table 1.

Long Ranger 2.2.2 (13) was used to align and call variants against GRCh38 for the 10XG NA12878 sample and was down-sampled (`-downsample 105`) from 75x coverage to approximately 30x coverage. GATK 4.0.0.0 (39) was used internally by Long Ranger to produce variant calls. RTG-Tools vcfeval 3.10.1 (40) was used to generate a VCF with the variants intersecting NA12878 Long Ranger 2.2.2 calls and the Platinum Genomes VCF. PacBio and Nanopore reads were aligned using minimap2 v2.11 (41) with recommended default settings for each respective technology. WhatsHap v0.18 (12) was to phase Platinum Genome variants (`whatshap phase`) with either PacBio or Nanopore reads at 20x coverage (`-max-coverage 20`) for read-selection, which included all variants and excluded read-groups for read selection (`-indels -ignore-read-groups`) with local realignment on (`-reference`) GRCh38 p.12. These steps produced a single VCF for each sequencing technology which incorporates phase calls. Visualization of phase blocks was completed using karyoploteR (42).

Each callset was then compared to the original Platinum Genomes VCF using whatshap compare to benchmark accuracy, where the error rate was averaged over all chromosomes. The `whatshap stats` command was used to generate phasing statistics for the four phased sets and GRCh38 p.12 chromosome lengths were provided (`-chr-lengths`) to calculate the phase block N50. A custom python script was used to assess each individually phased VCF to quantify phasing accuracy of adjacent heterozygous variants. The number of heterozygous variant pairs that were either unphased, phased correctly, or phased incorrectly was counted and benchmarked against the NA12878 platinum genome.

A Python script was written to combine phased VCF files generated from different technologies and output a VCF with a weighted consensus of the phase calls. For each adjacent heterozygous variant pair, a consensus call was generated by taking the most common phase configuration observed in the input VCF files. Each input VCF was weighted to allow ties to be broken. This script was used to generate a consensus phase callset for HG002 and the technologies included were weighted as follows: GIAB pedigree calls >Strand-Seq >10XG >PacBio CCS >PacBio CLR >Nanopore. This weighting scheme was based on the accuracy of these technologies. Python scripts can be found at (24)

### High molecular weight DNA extraction methods

Blood samples were extracted from patients with CF across Canada (Supplementary Table 6) and sent for processing to The Hospital for Sick Children in Toronto, Canada. Written informed consent was obtained from all participants, or parents/guardians/substitute decision makers. High molecular weight (HMW) DNA was extracted from fresh or frozen blood aliquots using the MagAttract HMW DNA Kit (Qiagen, Cat# 67563) as per supplier recommendations. Quantitation was determined by Quant-iT PicoGreen dsDNA Assay Kit (Invitrogen, Cat# P11469), as recommended by the supplier. Quality of DNA was then further assessed by electrophoretic migration in 0.4% agarose gel, run at 50 V for 18 hours at 4°C in Tris-acetate buffer at pH 8.0 with comparison to Quick-Load 1 kb Extend DNA ladder (NEB, Cat# N3239S). Unless otherwise stated, only samples indicating that bulk DNA was larger than 50 kb (>80% by visual inspection of agarose gel) were submitted for sequencing.

We also investigated three other DNA extraction methods including two Autopure methods (Maxi, 7-10 ml of blood; Midi, 3-4 ml of blood) and Puregene (0.3-1 ml of blood, manual extraction) (Qiagen, Cat# 1057048, 949006, 949008, 949016, 949018 and 949010). These samples were prepared as recommended by the kit supplier, but typically failed the HMW quality control assessment by the 0.4% agarose gel.

### Library preparation and 10x Genomics sequencing

Approximately 1 µg of genomic DNA was submitted to The Centre for Applied Genomics (TCAG) at the Hospital for Sick Children for genomic library preparation and whole genome sequencing. DNA samples were quantified using Qubit High Sensitivity Assay and sample purity was checked using Nanodrop OD260/280 ratio. DNA was run on the Genomic Tape on Tapestation (Agilent, Cat# 5067-5365 and 5067-5366) to check DNA fragment size. 10 ng of DNA was used as input material for library preparation using the 10XG Library Kit (PN 120258 and 120257) following the manufacturer’s recommended protocol. In brief, DNA was denatured and mixed with gel beads to form emulsion droplets using the Chromium Controller (PN 110203); emulsion droplets were tagged with barcodes and amplified by PCR; emulsions were broken and DNA captured and cleaned using magnetic beads. DNA was checked on the Bioanalyzer DNA High Sensitivity chip to ensure fragment size, and the DNA proceeds to library preparation. DNA was end-repaired, A-tailed, ligated with Illumina-compatible adapters, and PCR amplified with indexed Chromium i7 primers (PN 120262). Libraries are validated on a Bioanalyzer DNA High Sensitivity chip to check for size and absence of primer dimers and quantified by qPCR using Kapa Library Quantification Illumina/ABI Prism Kit protocol (KAPA Biosystems). Validated libraries were paired-end sequenced on an Illumina HiSeq X platform following Illumina’s recommended protocol to generate paired-end reads of 150-bases in length.

### Variant calling and phasing metrics for 10XG samples

Long Ranger 2.2.2 and GRCh38 reference version 2.1.0 were used process 10XG reads. Base calling was performed using the `mkfastq` command. VCF files were generated using the `wgs` command to call and phase variants; GATK 4.0.0.0 was used internally by Long Ranger to call variants. Alignment and phasing statistics were also generated by Long Ranger as output to the `wgs` command. The `stats` command from WhatsHap v0.18 was applied to the Long Ranger VCF files to produce additional phasing statistics. When both Long Ranger and WhatsHap reported the same metric, we took the values reported by Long Ranger. For causal CF variants, chart review and manual inspection of the Long Ranger alignment file with IGV was performed to investigate disagreements between clinical records and called variants.

### Generating genome graph from haplotypes

Using a multisample VCF of 449 10XG samples (all sequenced samples available at the time of analysis), variants were filtered to only include 50 bp surrounding exonic CFTR variants (GRCh38 chr7:117480087-117668359). Variants were further filtered to only include those with an rsID and of three or more. The intronic poly-T tract polymorphisms were manually called and phased using the 10XG sequencing reads. A graph representation of the haplotypes was generated using vg toolkit 1.33.0 (43) and plotted by Sequence Tube Map (44).

### 10XG Realignment and Deletion Polymorphism Calling

10XG sequencing reads aligned to the *PRSS1*-*PRSS2* locus (GRCh38 chr7:142500000-143000000) and a region spanning *PRSS3* (GRCh38 chr9:33700000-33900000) were extracted from the Long Ranger BAM file using SAMtools v1.9 (45). The extracted reads were realigned using Long Ranger 2.2.2 to a custom reference containing KI270803.1 and the *PRSS3* locus (GRCh38 chr9:33500000-34100000). The *PRSS3* locus was included because it shares a high base pair identity to the *PRSS1*-*PRSS2* locus, and we observed some reads aligned to *PRSS3* map better to the chromosome 7 locus.

To call the large deletion polymorphism observed on KI270803.1, a custom python script was used to determine the presence of the deletion by comparing the coverage of the deleted region (KI270803.1:771000-790000) to a flanking region of the same size (KI270803.1:760500-770000 and KI270803.1: 791000-800500) on both sides of the deleted region. Deletion calls were also visually validated using IGV. A dummy SNP was added to the VCF to encode the genotype of the deletion. An additional step was required to phase heterozygous deletion calls with respect to the other variants called by Long Ranger. Using haplotype-tagged 10XG linked reads, all heterozygous deletion calls were manually phased using IGV with respect to rs3757377 which lies upstream of the deletion. In the case where the deletion was heterozygous and rs3757377 was homozygous, the deletion was instead phased with respect to rs6666. Phase of the deletion calls in the VCF were updated using a custom script to reflect the phase relationship observed in the linked-reads.

Each 10XG VCF was filtered for variants with `PASS` in the `FILTER` column. Using bcftools 1.12 (46) merge, a multi-sample VCF was created by combining all the individual VCFs (`-missing-to-ref`). Variants in the multi-sample VCF called outside of KI270803.1 were removed. Variants with allele counts less than three, multi-allelic variants and indels longer than 5 bases (other than the 20 kb deletion which was coded as a SNP) were removed. SHAPEIT version 4.1.2 (22) was used to impute the missing variants and completely phase the multi-sample VCF to enable use as a reference panel (`-use-PS 0.0001 -sequencing`). Linkage disequilibrium blocks were computed from this VCF using ldblockshow version 1.36 (47) (`-BlockType 2 -SeleVar 1`).

### Illumina Genotype Arrays and Quality Control

CGMS data are genotyped on four different Illumina platform: 610Quad, 660W, Omni2.5 and Omni5. Genotype calling was performed using GenomeStudio V2011.1. Quality control steps were performed separately for each platform and described in detail in (10). Briefly, PLINK (48) was used for most QC steps while KING (49) identified any cryptic familial relationships among all individuals and PC-AiR (50) calculated PCs. Parents in six parent-offspring pairs, 19 samples clustered with Hapmap3 (51) African and East Asian ethnicity and 10 samples with sex-mismatch were excluded. Significant PCs were selected to be included in the association based on the Tracy-Widom test result using the function `twtable` in POPGEN of Eigensoft (52).

For colocalization of MI association with GTEx eQTLs, GWAS summary statistics (10) were reformatted as BED file and lifted to GRCh38 by LiftOver (53) for colocalization analysis against GTEx v8 in LocusFocus (29).

### Imputation of Genotype Data Using 10XG

Genotype array data was generated against GRCh37 and required lifting to alternative contig KI270803.1 before imputation. A two-step lift-over was performed using Picard LiftoverVcf (54); first from GRCh37 to GRCh38 using a chain file provided by UCSC and then from GRCh38 to alternative contig KI270803.1. The chain file from GRCh38 to KI270803.1 was created by downloading a PSL file for alternative haplotypes using the UCSC table browser and converting to a chain file using axtChain. Genotype array calls were organized by array platform into separate multi-sample VCF files and imputed by BEAGLE v5.1 (11) using the 10XG reference panel and default parameters.

### Association with MI

Variants from 2635 pancreatic insufficient individuals with BEAGLE imputation quality DR2 >0.3 were kept for association analysis with MI using imputation dosage of each variant, which was performed using the `geeglm` function from the R geepack package (55), with exchangeable correlation structure and binomial family. Sex, array platform and 11 PCs were included in the model. For conditional analysis, the dosage of the deletion was added as a covariate. For association testing with the 10XG data, only pancreatic insufficient individuals with available MI status were considered. 10XG variant calls within the range KI270803.1:700000-900000 were regressed against MI status (n=337 samples) using logistic regression. For conditioning on deletion genotype or rs62473563, the respective dosage was included as a covariate in the model. A subsequent regression was conducted where 28 individuals with the highest *CFTR* severity score were excluded.

### Re-processing of GTEx RNA-seq data

A custom reference genome was generated by adding the alternative contig KI270803.1 to a GRCh38 reference FASTA file. To remove sequence redundancy, the region on the chromosome 7 main contig corresponding to KI270803.1 (chr7:142038121-143088503) was masked with the ambiguous base “`N`”. 172 RNA-seq GTEx samples from pancreas were downloaded and reads were aligned to our custom reference using the scripts from the GTEx pipeline (56). First, GENCODE v26 (57) annotations were retrieved from the GTEx Portal and annotations within chr7:142038121-143088503 were removed. GENCODE v35 annotations for KI270803.1 were downloaded and collapsed using `collapse_annotation.py` available from the GTEx pipeline. The two resulting GTF files were combined into a single annotation file. We indexed our custom reference assembly with this annotation file using STAR v2.7.0 (58) (`-sjdbOverhang 75`). For each sample, we aligned RNA-seq reads using the `run_STAR.py` script from the GTEx pipeline. Transcript quantification was performed by mmquant (59) (`-l 20`) and read counts were normalized by conversion to transcripts per million (TPM).

### Recalculating GTEx Pancreas eQTL Data

Calculation of eQTLs was performed following the GTEx pipeline (56). GTEx v8 variant calls were filtered to chr7:142038121-143088503 and only included 252 pancreas samples with race labelled as “`white`”. Using the previously generated chain file, the GTEx multi-sample VCF and annotation BED file was lifted over from GRCh38 to KI270803.1. BEAGLE v5.1 was then used to impute the deletion from the 10XG reference panel into the GTEx VCF. Matching GTEx v8 read counts were normalized between samples using TMM (60). PEER factors were calculated from the normalized gene expression values using `run_PEER.R` from the GTEx pipeline. In addition to 15 PEER factors, the covariates used by GTEx v8 were included (five PCs, sex, PCR status and platform). FastQTL v2.184 (61) performed the eQTL analysis restricted to gene annotations on KI270803.1. For conditioning on deletion genotype or rs62473563, the respective dosage was included as a covariate in the model.

## Abbreviations

CF: cystic fibrosis
CFTR: cystic fibrosis transmembrane conductance
WGS: whole genome sequencing
GWAS: genome-wide association studies
MI: meconium ileus
LD: linkage disequilibrium
PacBio: Pacific Biosciences
10XG: 10x Genomics
CGMS: Canadian CF Gene Modifier Study Consortium
CLR: PacBio continuous long-reads
CCS: PacBio circular consensus sequence
VCF: variant call format
GIAB: Genome in a Bottle
HMW: high molecular weight
TCAG: The Centre for Applied Genomics
GTEx: Genotype-Tissue Expression
ER: endoplasmic reticulum
SRP: signal recognition particle
QC: quality control

## Ethics approval and consent to participate

The Canadian CF Gene Modifier Study (CGMS) was approved by the Research Ethics Board of the Hospital for Sick Children (#0020020214 from 2002-2019 and #1000016662 from 2019-present) and all participating sub-sites. Written informed consent was obtained from all participants or parents/guardians/substitute decision makers prior to inclusion in the study. The CGMS is approved by the Research Ethics Board of the Hospital for Sick Children for the usage of public and external data.

## Consents for publication

Not applicable.

## Availability of data and materials

The datasets generated and analyzed in this paper are publicly available. Data sources for NA12878 and HG002 reads and variant calls are summarized as Supplementary Table 1. Data from the CGMS analyzed for MI association including the genotype data are available from Canadian CF registry at https://www.cysticfibrosis.ca/our-programs/cf-registry/requesting-canadian-cf-registry-data. GTEx RNA-seq data and GTEx v8 variant calls were downloaded from dbGaP (accession number phs000424.v8.p) and the GTEx Portal https://www.gtexportal.org/home/datasets/, respectively.

## Code availability

All code and analyses steps implemented for phasing comparison with multiple sequencing techniques are available at https://github.com/strug-hub/cohort-phasing. Recalculation of GTEx eQTLs was performed following the GTEx pipeline: https://github.com/broadinstitute/gtex-pipeline.

## Acknowledgements

We thank the patients, care providers and clinic research assistants, collaborators, and principal investigators involved in CF Centers throughout Canada for their contributions to the CF Canada Patient Registry and Canadian Gene Modifier Study. The authors wish to acknowledge the staff supporting the High Performance Computing cluster and research helpdesk department and The Centre for Applied Genomics at the Hospital for Sick Children, Toronto. The Genotype-Tissue Expression (GTEx) Project was supported by the Common Fund of the Office of the Director of the National Institutes of Health, and by NCI, NHGRI, NHLBI, NIDA, NIMH, and NINDS.

